# Lineage-specific dynamics of erasure of X-upregulation during inactive-X reactivation

**DOI:** 10.1101/2020.12.23.424181

**Authors:** HC Naik, D Chandel, S Majumdar, M Arava, R Baro, H Bv, K Hari, Parichitran, Avinchal, MK Jolly, S Gayen

**Affiliations:** Department of Development Biology and Genetics, Indian Institute of Science, Bangalore-560012, India; Centre for BioSystems Science and Engineering, Indian Institute of Science, Bangalore 560012, India; IISc Mathematics Initiative (IMI), Indian Institute of Science, Bangalore 560012, India

**Keywords:** X-chromosome inactivation, iPSC, Primordial germ cells (PGCs), Pre-implantation, pre-gastrulation, epigenetics, B-cells, extra-embryonic endoderm stem (XEN), Epiblast stem cells (EpiSC)

## Abstract

In mammals, sex chromosome dosage is compensated through X-chromosome inactivation and active-X upregulation. It is believed that during early development, X-chromosome inactivation and active X upregulation happen in a highly coordinated fashion. However, such coordination between two X-chromosomes in other developmental contexts remains unexplored. Here, we have profiled the coordination between two X-chromosomes in female cells in different developmental contexts and cell types: pre-implantation embryos, embryonic epiblast cells, iPSC reprogramming, germ cell reprogramming, B-cell, and extra-embryonic endoderm stem (XEN) cells. Interestingly, we found that two X-chromosomes in female cells are not always coordinated; instead, it happens in a lineage-specific manner. Specially, while embryonic mouse epiblast cells, iPSC undergo erasure of X-upregulation upon reactivation of the inactive X, germ cells do not. Importantly, we show that the erasure of X-upregulation in epiblast or iPSC is potentially mediated via undifferentiated embryonic transcription Factor 1 (UTF1), which is absent or lowly expressed in late germ cells and therefore, germ cells are unable to erase upregulation. Moreover, we found that partial reactivation of the inactive X is insufficient to drive the erasure of upregulation globally, nor from their counterparts on the active X in XEN and B-cells. Finally, through a phenomenological mathematical model, we show that cross-inhibition between two X-chromosomes can reproduce the dynamics of reactivation and erasure of upregulation. Altogether, our study provides insight into the coordination between two X-chromosomes in female cells in different developmental contexts and related mechanistic aspects.

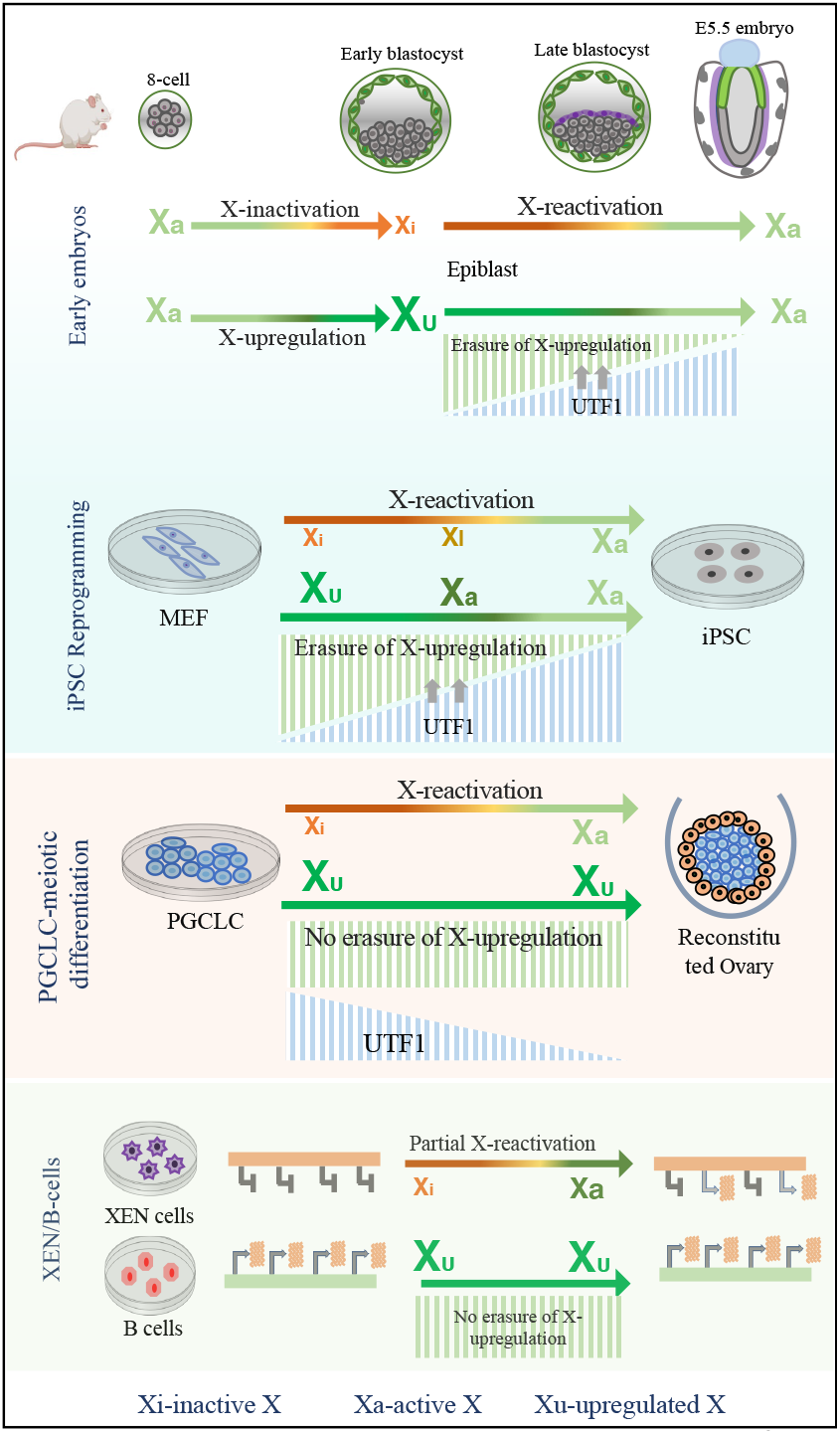

## Introduction

In therian female mammals, one of the X-chromosomes in females is inactivated to compensate the dosage of X-linked gene expression between the sexes (Gayen et al., 2016; Lyon, 1961; Samanta et al., 2022). On the other hand, male has only one X-chromosome. Therefore, X-linked genes are monoallelically expressed in both sexes compared to the biallelic expression of autosomal genes. Proper dosage of X-linked gene expression relative to autosomal genes is crucial to drive precise development and cell fate specification. A longstanding question is how the dosage of monoallelic X-linked genes is compensated relative to the autosomal genes. The upregulation of X-linked genes from the active-X chromosomes in both sexes is thought to balance the X-linked gene dosage relative to autosomal genes. The concept of X-chromosome upregulation was first hypothesized by Ohno in 1967 (Ohno S, 1967). As per Ohno, during the evolution of sex chromosomes from a pair of autosomes, the Y chromosome’s degradation led to the dosage imbalance between X-linked and autosomal genes in males, which was balanced through the upregulation of X-linked genes (Graves, 2016). Subsequently, to counteract the extra dosage from the X chromosome in female cells, the evolution of X-chromosome inactivation happened. While X-inactivation has been extensively studied, the existence of active-X upregulation remains controversial and has been challenged for a long time (Chen and Zhang, 2016; Deng et al., 2011; Julien et al., 2012; Kaur et al., 2020; Kharchenko et al., 2011; Xiong et al., 2010). Recent studies by us and others have provided substantial shreds of evidence for the existence of upregulated active-X chromosome in both male and female cells in mouse and human (Deng et al., 2013; Larsson et al., 2019; Lentini et al., 2022; Li et al., 2017; Lin et al., 2011; Mandal et al., 2020; De Mello et al., 2017; Naik et al., 2022; Sangrithi et al., 2017). However, the developmental dynamics of X-upregulation have not been explored much. Recently, through single-cell analysis, we have demonstrated that X-upregulation happens concomitantly upon initiation of X-inactivation in pre-gastrulating mouse embryos (Naik et al., 2022). Overall, it is believed that two X-chromosomes in female cells coordinate with each other to fine-tune the X-linked gene expression during early development. Whether such coordination between two X-chromosomes in female cells is a universal phenomenon or restricted to certain developmental windows or cell types remains unknown. Specially, the transcriptional kinetics of two X-chromosomes during the reactivation of inactive-X genes in different developmental contexts, such as in mouse embryonic epiblast, iPSC reprogramming and germ cell reprogramming, remains poorly understood. To address these, we have profiled the coordination between two X-chromosomes in female cells in different developmental contexts and cell types: pre-implantation embryos, embryonic epiblast cells, iPSC reprogramming, germ cell reprogramming, B-cell, and extra-embryonic endoderm stem cells (XEN). Interestingly, we found that two X-chromosomes in female cells are not always coordinated; instead, it happens in a lineage-specific manner. Specially, we show that while embryonic mouse epiblast cells, iPSC undergo erasure of X-upregulation upon reactivation of the inactive X, germ cells do not. On the other hand, mechanistic aspects of X-chromosome upregulation/erasure of X-upregulation and how two X-chromosomes in female cells coordinate with each other remain poorly understood. It is thought that increased transcriptional burst frequency and enrichment of different active chromatin marks on the active-X drive the X-upregulation (Larsson et al., 2019; Naik et al., 2022). However, what drives the erasure of X-upregulation during the reactivation of the inactive-X is not known yet. Here, we have identified undifferentiated embryonic transcription Factor 1 (UTF1) as the potential key regulator for the erasure of X-upregulation. Taken together, our study provides significant insight into the expression dynamics of two X-chromosomes in different developmental contexts and related mechanistic aspects.

## Result

### Two X-chromosomes coordinate with each other to fine-tune X-linked gene expression in female pre-implantation embryo

In mice, X-inactivation happens in two phases: initially imprinted X-inactivation in pre-implantation embryos, where paternal-X gets inactivated and later switches to random X-inactivation in the post-implantation epiblast (Harris et al., 2019; Maclary et al., 2017; Saiba et al., 2018). Switching of imprinted to random X-inactivation is mediated through the reactivation of the imprinted inactive-X in the epiblast cells (Gayen et al., 2015; Maclary et al., 2014; Sarkar et al., 2015). To understand the active-X expression dynamics during initiation of imprinted X-inactivation during pre-implantation development, we have profiled allelic X-linked gene expression at different stages of mouse pre-implantation embryos (8-cell, 16-cell, 32-cell, E3.5 early, E4.0 mid and E4.5 late blastocyst) using available single-cell RNA-seq (scRNA-seq) datasets (Fig. 1A). These embryos harbored polymorphic sites across their genome as they were derived from two diverged mouse strain: C57BL/6J and CAST, which allowed us to perform allele-specific gene expression analysis. First, we segregated cells of female embryos based on their X-chromosome inactivation status by profiling the fraction of maternal allele expression (Fig. 1B). As expected, we found three different categories of cells: cells with no X-inactivation (XaXa), partial X-inactivation (XaXp), and complete X-inactivation (XaXi), indicating cells are on set to establish imprinted X-inactivation (Fig. 1B). In case of male cells, we found expression of X-linked genes from maternal allele only (Fig. 1B). Additionally, autosomal genes had equivalent allelic expression from the paternal and maternal allele, thereby validating our allele-specific expression analysis methodology (Fig. S1A). Next, we profiled allelic X to autosomal (X_mat_:A_mat_ and X_pat_:A_pat_) gene expression ratio in different categories of cells (XaXa, XaXp and XaXi) of female embryos to delineate the active-X upregulation dynamics upon initiation of X-inactivation. X:A ratio is used as an indicator for the presence of X-upregulation. If there is upregulation from maternal-X, the expected X_mat_:A_mat_ ratio should be >1 and close to 2. We found that X_mat_:A_mat_ and X_pat_:A_pat_ ratio in XaXa cells was almost equivalent and slightly greater than 1, indicating overall higher X-linked expression than the autosomal genes during pre-implantation development (Fig 1C). As expected, we found that in XaXp cells, X_pat_:A_pat_ ratio tends to be down and reaches almost none in XaXi cells, indicating inactivation of the paternal-X (Fig 1C). Interestingly, in XaXp and XaXi cells, we found that active-X_mat_:A_mat_ ratio dynamically increased to ~1.5-2, indicating dynamic X-upregulation upon X-inactivation (Fig 1C). Similarly, in male cells, we found that X_mat_:A_mat_ ratio is > 1 and close to 2, suggesting active-X upregulation in male pre-implantation embryos (Fig. 1C). Next, to get more insight into the expression dynamics of X-linked genes, we profiled the allelic expression pattern from autosome and X-chromosome. Interestingly, we found that active X expression in female XaXp/XaXi cells and male cells is always higher compared to the autosomal allelic expression, corroborating the upregulation of gene expression from the active X-chromosome (Fig. 1C). Next, to get insight into the developmental stage-wise dynamics of two X-chromosomes, we performed a similar analysis across different stages of mouse pre-implantation embryos (8-cell, 16-cell, 32-cell, E3.5 early, E4.0 mid and E4.5 late blastocyst) in XaXa, XaXp and XaXi cells. We found a similar pattern that upon X-inactivation, there is dynamic X-upregulation of the maternal X chromosome beginning from 8-cell stage (Fig. 1D). Similar pattern was also found in male cells (Fig. 1D). Altogether, this analysis suggested that upon X-inactivation in mouse pre-implantation embryos, concomitant active X-upregulation fine-tune the X-linked gene dosage relative to autosomal genes.

**Figure 1:**
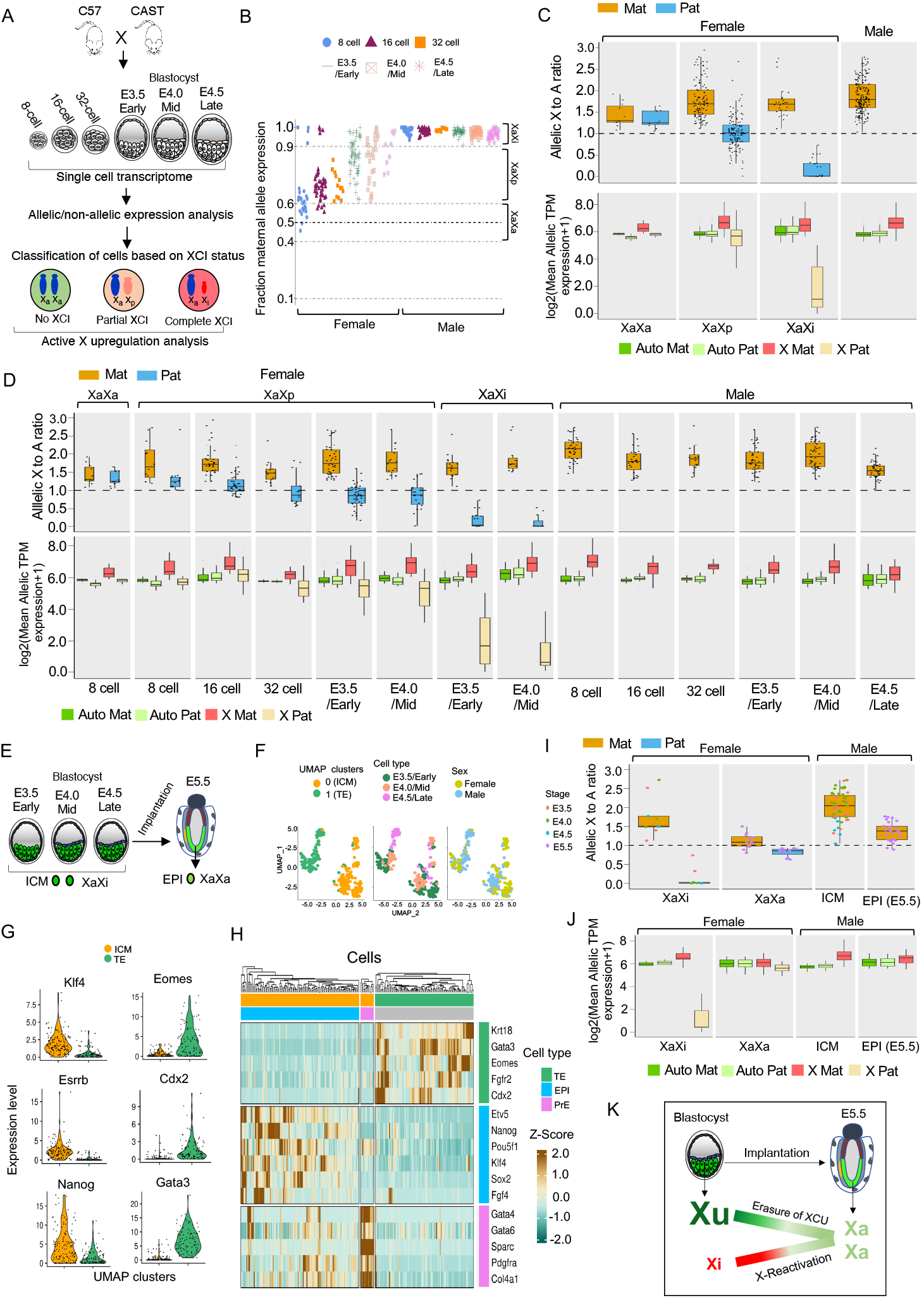
Erasure of active-X upregulation upon reactivation of inactive-X in the embryonic epiblast. (A) Schematic representing experimental designing of profiling active X upregulation dynamics throughout different stages of mouse pre-implantation hybrid embryos (8-cell, 16-cell, 32-cell, E3.5 early, E4.0 mid and E4.5 late blastocyst) at the single-cell level using scRNA-seq dataset. These embryos were generated from the crossing of two divergent mouse strains, C57 and CAST (B) Classification of female cells of pre-implantation embryos based on X-inactivation state through profiling of fraction maternal expression of X-linked genes. Male cells showed expression from maternal allele only (C) Allelic X:A ratio and allelic expression (log mean allelic TPM+1) plots for the XaXa, XaXp, XaXi female cells and male cells of pre-implantation embryos. (D) Allelic X:A ratio and allelic expression (log mean allelic TPM+1) plot for the XaXa, XaXp, XaXi female cells and male cells throughout different stages of pre-implantation development. (E) Schematic showing embryonic epiblast cells of preimplantation and post-implantation (E5.5) embryos. (F) UMAP clustering of epiblast (inner cell mass) and TE cells of pre-implantation embryos (E3.5 early, E4.0 mid and E4.5 late blastocyst). (G) Violin plots showing the expression of different markers corresponding to EPI and TE cells in UMAP-based clusters. (H) Heatmap representing the expression of markers of EPI, TE and PrE clusters. (I) Comparison of allelic X:A ratio between XaXi and XaXa epiblast female cells and between ICM and post-implantation male epiblast cells. (J) Allelic expression (log mean allelic TPM+1) of X-linked and autosomal genes for XaXi and XaXa female and male epiblast cells. (K) Model showing erasure of active X upregulation upon reactivation of inactive-X in embryonic epiblast cells.

### Erasure of active-X upregulation upon reactivation of inactive X in the embryonic epiblast

Next, we interrogated if X-upregulation is erased during the reactivation of the inactive-X chromosome in epiblast cells transitioning from imprinted to random X-inactivation. To check this, first, we identified inner cell mass (ICM) or epiblast precursor cells from E3.5, E 4.0 and E4.5 cells through UMAP clustering and marker expression (Fig. 1E, F, G and H). We found two major clusters corresponding to ICM expressing ESRRB, NANOG, KLF4 and trophectoderm (TE) showing expression of EOMES, GATA3, CDX2 (Fig. F and G). Further, we segregated into three different lineages of the blastocyst: epiblast (EPI), primitive endoderm (PrE) and TE based on marker gene expression clustering (Fig. 1H). As there were few XaXa cells in the late blastocyst, we used E5.5 XaXa epiblast cells as identified in our previous study (Naik et al., 2022). Next, we compared the allelic X:A ratio between XaXi vs. XaXa cells of epiblast (Fig. 1I). Notably, we found that X_mat_:A_mat_ ratio in reactivated (XaXa) epiblast cells reduced significantly compared to the XaXi cells indicating the erasure of upregulation in these cells (Fig. 1I). This notion was further supported by allelic expression from maternal-X in these cells, which showed no significant difference with the autosomal allelic expression (Fig. 1J). Taken together, these analyses suggested the erasure of X-upregulation in female epiblast upon reactivation of inactive-X (Fig. 1K). We extended our analysis to male cells as well. We compared the X_mat_:A_mat_ ratio of pre-implantation male epiblast (ICM) cells with post-implantation epiblast cells and found that there is a reduction in X_mat_:A_mat_ ratio in post-implantation epiblast cells, albeit X:A ratio was still close to 1.5 (Fig. 1I). Further, allelic analysis confirmed presence of upregulated X in the post-implantation male epiblast (Fig. 1J). Overall, may be at this stage of development male cells undergo transient erasure of X-upregulation.

### Dynamic erasure of active-X upregulation upon reactivation of inactive-X during iPSCs reprogramming

Next, we investigated the active-X expression dynamics during the reactivation of the inactive X at different stages of the reprogramming of female mouse embryonic fibroblast (MEF) to induced pluripotent stem cells (iPSC). We used the available bulk RNA-seq dataset of Oct4, Sox2, Klf4, and Myc (OSKM) mediated iPSC reprogramming from Janiszewski et al. (Janiszewski et al., 2019) (Fig. 2A). MEF cell line used for the reprogramming in their study was derived from two divergent mouse strains: Mus musculus musculus (mus) and Mus musculus castaneous (cast) and therefore harbored polymorphic sites between the paternal and maternal chromosomes, which allowed us to do allele-specific expression analysis of X-linked and autosomal genes throughout the different stages of reprogramming (Fig. 2A). Notably, X-inactivation is skewed to the mus allele in these cells, which helped to distinguish allelic X-linked gene expression from the active vs. inactive X-chromosome. We analyzed d2 MEF, different intermediate FUT4 positive cells (d8, d10, d13 and d15) and iPSC (Fig. 2A). First, we profiled allelic X:A ratio in these cells. We found that X_cast_:A_cast_ ratio in d2 MEF was close to 2, indicating the presence of upregulated active X-chromosome. Intriguingly, we found a decrease in X_cast_:A_cast_ ratio from d10 onwards upon increase in X_mus_:A_mus_ ratio, suggesting concomitant erasure of active-X upregulation upon X-reactivation (Fig. 2B). To probe this further, we delineated the dynamics of erasure of X upregulation of reactivated X-linked genes throughout the different stages of reprogramming. We identified reactivated X-linked genes throughout the different stages of reprogramming by profiling the fraction of allelic expression from the inactive-X allele (mus) (Fig. 2C). We considered a gene as reactivated if it showed fraction of allelic expression from the inactive-X allele above 0.1 (Fig. 2C). We found indeed many X-linked genes showing the decrease in expression from the active X_cast_ chromosome upon reprogramming (Fig. 2D). Altogether, these analyses indicated erasure of X-chromosome upregulation along with reactivation of the inactive X chromosome during iPSC reprogramming. Next, to get better insight into the coordination of two X-chromosomes during reprogramming, we profiled the dynamics of two X chromosomes during iPSC reprogramming at single cell level through allele-specific analysis of scRNA-Seq dataset from Janiszewski et al. (Janiszewski et al., 2019). MEF cells used for the reprogramming was hybrid (cast X mus) and inactivation is skewed towards the mus allele as described above, which enabled us to profile allelic gene expression and distinguish expression from active vs inactive X (Fig. 2E). We analyzed d0 MEF, different intermediate SSEA1 positive cells (d8, d9, d10, d12) and iPSC (Fig. 2E). First, we presumably categorized cells into four different categories based on their extent of reactivation through profiling fraction of allelic expression of X-chromosomes: cells with inactive X (Xi), cells with partial reactivation (Xr-Low), intermediate reactivation (Xr-intermediate) and robustly reactivated (Xr-Robust) (Fig. 2F). As expected, we found while d0 MEF cells belonged to Xi category, iPSC belonged to Xr-Robust or Xr-intermediate categories (Fig. 2F, Fig. S2). Cells of d8, d9 and d10 belonged majority to Xi cells along with few Xr-Low and Xr-intermediate cells, whereas many cells of d12 belonged to Xr-robust or Xr-intermediate (Fig. 2F, Fig. S2). Next, we profiled allelic X:A ratio in these different categories of cells to delineate the correlation between extend of reactivation vs erasure of X-upregulation. We found that active-X_cast_:A_cast_ ratio in Xi cells is >1.5 and close to 2, indicating these cells harbor upregulated active-X chromosome (Fig. 2G). On the other hand, Xr-Low cells had almost similar active-X_cast_:A_cast_ ratio to Xi cells, indicating no erasure of X-upregulation (Fig. 2G). Whereas in Xr-robust or Xr-intermediate cells active-X_cast_:A_cast_ ratio reduced to 1, suggesting the erasure of X-upregulation in these cells (Fig. 2G). Allelic expression analysis corroborated a similar pattern (Fig. 2H). We performed a similar analysis of these different categories of cells through different days of reprogramming to disintegrate the reactivation kinetics vs. stages of reprogramming contributing to the erasure of X-upregulation (Fig. 2I). We found that there was no variation in the pattern of erasure of X-upregulation along with the stages of reprogramming instead it correlated with the status of X-reactivation, i.e., the erasure was triggered upon intermediate/robust reactivation of the inactive-X, suggesting that the erasure of X-upregulation is triggered after a certain threshold of reactivation during iPSC reprogramming (Fig. 2I). Altogether, our analysis suggested dynamic erasure of X-upregulation during the reactivation of the inactive-X during iPSC reprogramming (Fig. 2J).

**Figure 2:**
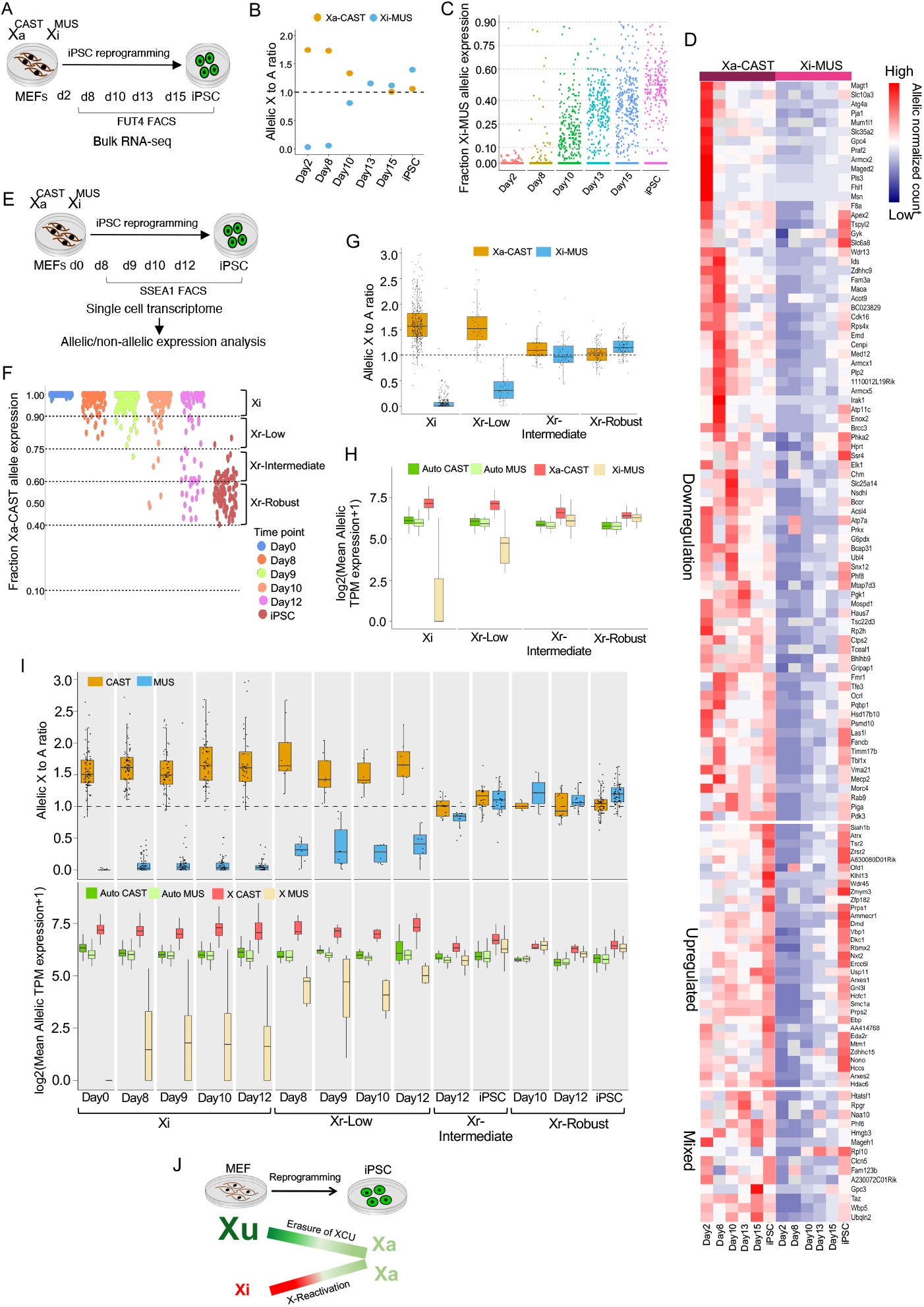
Dynamic erasure of active-X upregulation upon reactivation of inactive-X during iPSCs reprogramming. (A) Schematic showing different stages of reprogramming of female MEF to iPSC, in which allele-specific RNA-seq analysis was performed to assay X-upregulation dynamics using bulk RNA-seq dataset. (B) Allelic X:A ratio throughout the different stages of iPSC reprogramming. (C) Schematic of iPSC reprogramming stages along with X-reactivation status. (C) Plot representing fraction Xi-*Mus* allele expression throughout the different stages of iPSC reprogramming. (D) Heatmap representing allelic expression from Xi-*Mus* and Xa-*CAST* allele (Log2 normalized allelic reads) of X-linked genes throughout the reprogramming stages. Genes are classified into different categories based on the erasure kinetics during the reprogramming process. (E) Schematic showing different stages of reprogramming of female MEF to iPSC, in which allele-specific RNA-seq analysis was performed using single-cell transcriptomic dataset to assay X-upregulation dynamics. (F) Identification of different categories of cells based on X-reactivation status through profiling fraction Xa-*CAST*allele expression. (G) Allelic X:A ratio plot of different categories of cells: Xi, Xr-Low, Xr-intermediate and Xr-robust. (H) Allelic expression (log mean allelic TPM+1) of X-linked and autosomal genes in different categories of cells: Xi, Xr-Low, Xr-intermediate and Xr-robust. (I) Allelic X:A ratio and allelic expression (log mean allelic TPM+1) plot for Xi, Xr-Low, Xr-intermediate and Xr-robust cells throughout different stages of reprogramming. (J) Model showing dynamic erasure of active X upregulation upon reactivation of inactive-X during reprogramming of female MEF to iPSC.

### No erasure of active X upregulation upon reactivation of inactive X in germ cells

Germ cells are specified as primordial germ cells (PGCs) from the post-implantation epiblast of mouse embryo (Hajkova et al., 2002, 2008; Lawson et al., 1999). PGCs then migrate to the gonad, where they undergo sexual differentiation. During the migration to the gonad, PGCs undergo extensive epigenetic reprogramming to erase the parental information and establish new marks during gametogenesis (Hill et al., 2018). One such crucial epigenetic reprogramming in female PGCs is the reactivation of the inactive X chromosome (Sugimoto and Abe, 2007). It is thought that X-reactivation in female PGCs occurs in a stepwise manner during the migration to the gonad and gets completed at the gonad when they enter meiosis (Chuva De Sousa Lopes et al., 2008; Sangrithi et al., 2017). However, whether the reactivation of inactive-X in germ cells leads to the erasure of active-X upregulation remains poorly understood. Here, we have profiled the active-X expression dynamics during the reactivation of the inactive X in germ cells at the single-cell level. We have used the available scRNA-seq dataset of an *in vitro* germ cell differentiation system from Severino et al. (Severino et al., 2022). In brief, Severino et al. derived primordial germ cell like cells (PGCLC) from embryonic stem cells (ESC) and differentiated PGCLCs to facilitate their meiotic entry using an *in vitro* reconstituted Ovary (rOvary), which mimics female urogenital environment. We analyzed scRNA-seq data of differentiated germ cells, which originated from XGFP-negative PGCLC population as described in Severino et al. (Fig. 3A). XGFP-negative PGCLC population harbored inactive-X and underwent reactivation upon rOvary mediated differentiation and thereby served as a good system to track the erasure of X-upregulation upon X-reactivation. Additionally, these cells harbored *Mus musculus* (X^mus^) and *Mus castaneous* (X^cast^) X chromosomes, thereby allowed us to profile allele-specific X-linked gene expression analysis. Notably, X^mus^ is inactivated in these cells, and therefore allowing us to track down allelic X-linked gene expression from inactive-X vs. active-X. First, we segregated XGFP-PGCLC differentiated cells into mitotic and meiotic populations based on UMAP and marker-based clustering, as described by Severino et al. (Fig. 3 B). In consistence with Severino et al., we identified two different UMAP clusters representing mitotic and meiotic cells (Fig. 3 B). Finally, we have identified different stages of germ cell maturation: mitotic (mitotic 1 and 2) to early meiotic (pre-meiotic-1 and 2) and late meiotic germ cells based on marker gene expression-based clustering (Fig. 3B and 3C). Further, we categorized all these cells presumably into four different categories based on their extent of reactivation through profiling fraction of allelic expression of X-chromosomes: cells with inactive X (Xi), cells with partial reactivation (Xr-Low), intermediate reactivation (Xr-intermediate) and robustly reactivated (Xr-robust) (Fig. 3D). We found that majority of mitotic cells had robust X-reactivation (Fig. 3D and E). On the other hand, pre-meiotic and meiotic cells also had reactivation of the inactive-X, but most cells belonged to the Xr-intermediate category (Fig. 3D and E). To delineate if the reactivation of X-linked genes leads to the erasure of X-upregulation in germ cells, we profiled allelic X:A (Chr 13) ratio in these different categories of cells. As expected, active-X_cast_:A_cast_ ratio of Xi cells was >2, suggesting these cells harbor upregulated active X chromosome (Fig. 3F). Surprisingly, we found that cells with reactivated X-chromosome still had active-X_cast_:A_cast_ ratio >1.5 and close to 2, indicating there is no erasure of X-upregulation upon X-reactivation (Fig. 3F). Allelic expression analysis showed a similar pattern (Fig. 3G). Next, we analyzed allelic X:A ratio in mitotic, pre-meiotic and meiotic cells to identify if there is a stage-specific pattern of two X-chromosomes states. We found that mitotic cells had no erasure of X-upregulation, and surprisingly there was hyperactivation of the inactive-X, which became equivalent to the active-X expression (Fig. 3H). Pre-meiotic and meiotic reactivated cells still had upregulated active-X chromosome, but there was no hyper activation of the reactivated X-chromosome (Fig. 3H). Altogether, this analysis suggested that the reactivation of inactive-X in germ cells is not associated with the erasure of X-upregulation (Fig. 3I).

**Figure 3:**
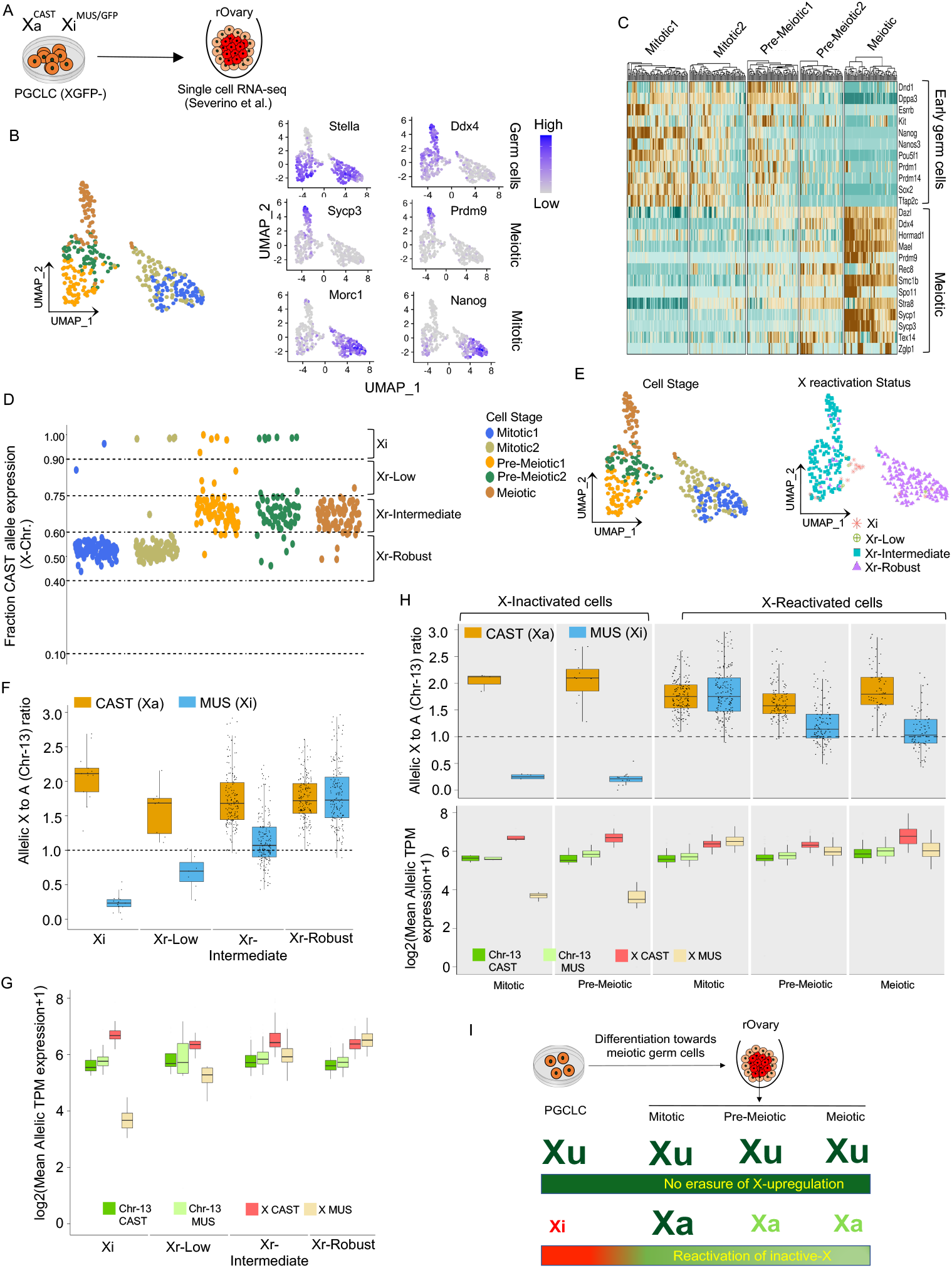
Active X-upregulation is not erased upon reactivation of inactive-X in germ cells. (A) Schematic showing differentiation of XGFP-PGCLCs towards meiotic cells using rOvary system as described in Severino et al. 2022. (B) Plots representing UMAP-based clustering and projection of marker gene expression on UMAP plot for mitotic and meiotic germ cells. (C) Heatmap based on gene expression dynamics representing different stages of germ cell maturation: mitotic 1, mitotic 2, early meiotic (pre-meiotic-1 and 2) and late meiotic germ cells. (D) Plot representing fraction Xa-*CAST* allele expression throughout the different stages of germ cell maturation. (E) Plot representing projection of X-reactivation status of cells on UMAP. (F) Allelic X:A ratio plot of different categories of cells: Xi, Xr-Low, Xr-Intermediate and Xr-Robust. (G) Allelic expression (log mean allelic TPM+1) of X-linked and autosomal genes in different categories of cells: Xi, Xr-Low, Xr-Intermediate and Xr-Robust. (H) Allelic X:A ratio and allelic expression (log mean allelic TPM+1) plot for cells with inactivated X and cells with reactivated inactive-X throughout different stages of germ cell maturation. (I) Model showing no erasure of active X upregulation upon reactivation of inactive X during germ cell reprogramming.

### Partial reactivation of the inactive-X does not trigger the erasure of X-upregulation in XEN cells

So far, in above experiments, we have analyzed the erasure of the active-X upregulation in the context of spontaneous X-reactivation in different lineages such as embryonic epiblast, iPSC and germ cells. Next, we tested further in a scenario where inactive-X reactivation does not occur naturally; instead, driven through some manipulation also trigger the ersure of X-upregulation or not. To test this, the primary technical limitation is that no method is available to fully reactivated the inactive-X in cells which normally maintain the inactivation state of the X-chromosome. However, inactive-X can be partially reactivated through the ablation of Xist. Xist is the master regulator of the X-inactivation, which coats the entire inactive-X and recruits different chromatin modifiers to facilitate transcriptional silencing of the inactive-X. Therefore, we attempted to ablate Xist in XEN cells to reactivate the inactive-X. XEN cells used for this study were derived from two divergent mouse strains, *Mus Musculus* (mus) and *Mus Molassinus* (mol), allowing us to perform allele-specific gene expression analysis. Importantly, X-inactivation is skewed towards paternal-X (mus) as XEN cells undergo imprinted X-inactivation, which helped to differentiate the X-linked gene expression from active-X vs. inactive X through allele-specific analysis (Fig. 4A). We ablated Xist in XEN cells using CRISPR-Cas9 based approach using two gRNAs targeting to Xist upstream region and intron1 respectively (Fig. 4A). To identify the clones with Xist ablation (sgXist), we performed RNA-FISH against Xist using fluroescently labeled Xist probe. We selected a clone for this study, which did not show the presence of coating of Xist in RNA FISH (Fig. 4B). RNA-Seq analysis further validated the complete lack of Xist expression in this sgXist cell line (Fig. 4C). Next, we estimated allelic X:A ratio in WT and sgRNA XEN cells. As expected, we found that active-X_mol_:A_mol_ ratio in WT XEN cells is close to 1, indicating presence of upregulated active-X chromosome. Similarly, inactive-X_mus_:A_mus_ ratio in WT XEN cells was close to zero, indicating inactivation of the mus X chromosome (Fig. 4D). On the other hand, we found that in sgXist XEN cells inactive-X_mus_:A_mus_ ratio increased to ~0.5, suggesting partial reactivation of the inactive X-chromosome (Fig. 4D). However, apparently there was no erasure of active-X upregulation as revealed by active-X_mol_:A_mol_ ratio in sgXist cells, which was similar to the WT XEN cells (Fig. 4D). Allelic expression analysis of X-linked and autosomal genes corroborated the similar facts (Fig. 4E). Next, we interrogated if there was erasure of upregulation of the X-linked genes, which were reactivated. To test this, first, we identified different categories of reactivated genes (Xr-Low, Xr-intermediate, Xr-robust) through fraction allelic expression analysis as described above (Fig. 4F). Next, we profiled allelic X:A ratio for the reactivated genes only. In WT XEN cells, active-X_mol_:A_mol_ ratio for reactivated genes was also close to 1.5, indicating these fractions of genes undergo upregulation (Fig. 4G). However, there was no change in active-X_mol_:A_mol_ ratio for reactivated genes in sgXist XEN cells indicating no erasure of X-upregulation from the counterpart of the reactivated genes on the active-X (Fig. 4G). Allelic expression analysis of reactivated X-linked genes and autosomal genes also showed higher expression of X-linked genes from active-X_mol_ allele compared to the autosomal allelic expression both in WT and sgXist XEN cells, thereby further confirming no erasure of X-upregulation from the cohort of reactivated genes (Fig. 4H). Finally, we profiled the gene-wise allelic expression of reactivated genes in WT and sgXist XEN cells, which also revealed not much change in expression from the active-X expression upon reactivation (Fig. 4I). Taken together, our analysis revealed that partial reactivation of the inactive-X in sgXist XEN cells neither leads to erasure of global X-upregulation nor from the counterpart of the reactivated genes.

**Figure 4:**
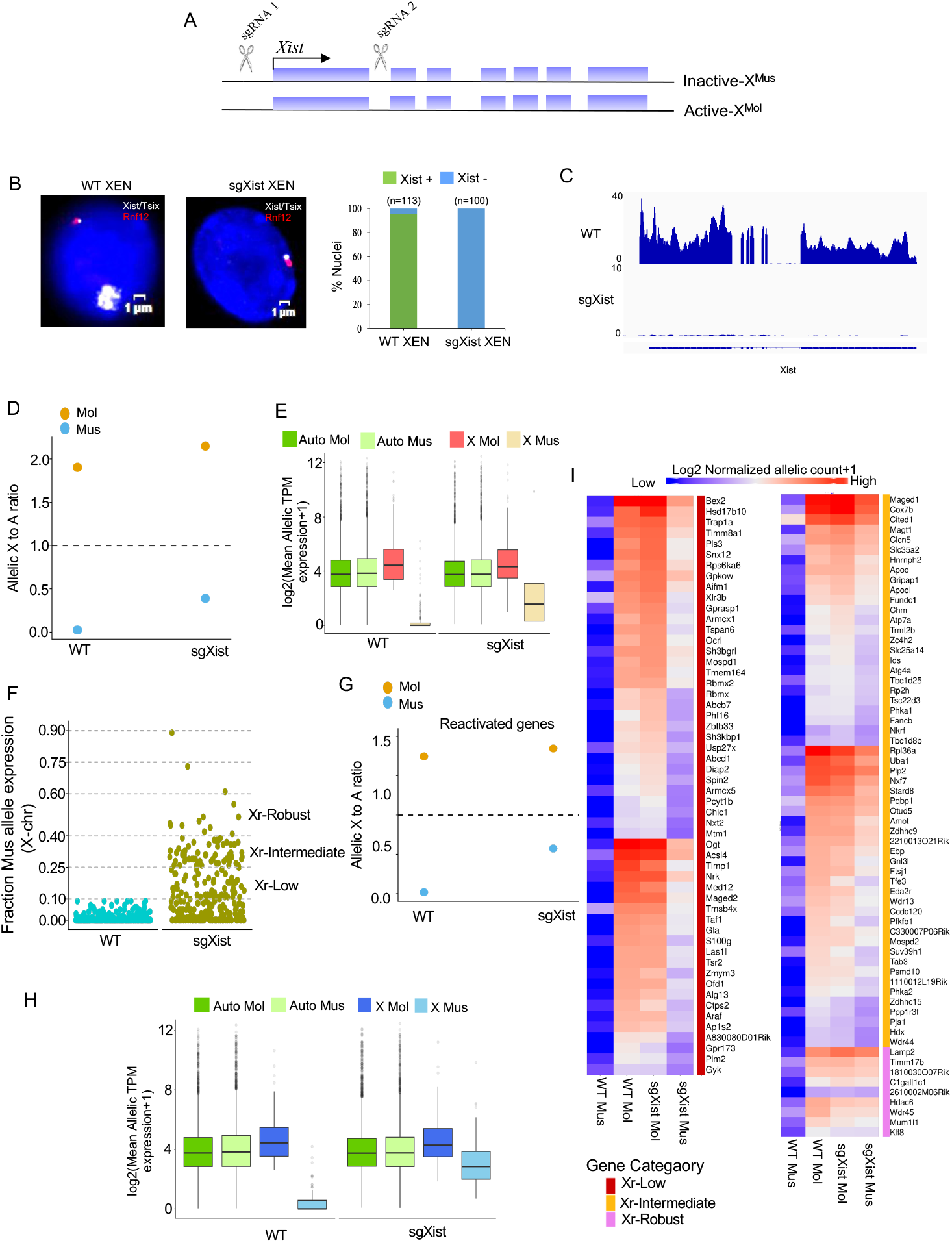
Partial reactivation of inactive-X does not lead to the erasure of active X-upregulation in XEN cells. (A) Schematic of sgRNAs targeting the Xist locus (B) Detection of Xist/Tsix RNA (white) and Rnf12 (red) through RNA-FISH in WT and sgXist XEN cells. DAPI stained the nuclei in blue. The scale bar represents 1 um. Right: quantification of Xist coated nuclei in WT and sgXist XEN cells. (C) RNA-seq signals for Xist in WT and sgXist XEN cells. (D) Plot representing allelic X:A ratio in WT and sgXist XEN cells. (E) Plot representing autosomal and X-chromosome allelic expression (Log mean allelic TPM+1). (F) Plot representing fraction paternal allele (Xi-*Mus*) expression in WT and sgXist XEN cells. (G) Allelic X:A ratio for reactivated X-linked genes. (H) Allelic expression (log mean allelic TPM+1) plot for autosomal and reactivated X-linked genes. (I) Heatmap representing allelic expression from Xi-*Mus* and Xa-Mol allele (Log2 normalized allelic reads) of X-linked genes.

### Partial reactivation of the inactive-X does not trigger the erasure of X-upregulation in human B-cells

Next, we extended our analysis to human B-cells having partially reactivated inactive-X from available RNA-Seq data (Yu et al., 2021). In these cells, XIST was knockdown through CRISPR interference (CRISPRi) of XIST (sgXIST). B-cells and sgXIST B-cells were also treated with DNMT and EZH2 inhibitors to ablate DNA methylation and H3K27me3. Further, these cells have skewed X-inactivation towards paternal-X chromosome and therefore allowed us to differentiate the X-linked gene expression from active vs. inactive-X. First, we performed allelic X:A ratio analysis in these cells and found that control B-cells harbor upregulated active-X chromosome as indicated by high (>2) active-X:A ratio (Fig. 5A). On the other hand, in sgXIST, inhibitors-treated and sgXIST+inhibitors cells, there was slight increase inactive-X:A ratio indicating partial reactivation of the inactive-X chromosome (Fig. 5A). However, there was no significant changes in active-X:A ratio in these cells suggesting no erasure of X-upregulation (Fig. 5A). Allelic expression of X-linked and autosomal genes corroborated the similar facts (Fig. 5B). Next, we extended our analysis to gene-wise manner focusing only the cohort of reactivated genes. As described above, we identified different categories of reactivated genes (Xr-Low, Xr-intermediate, Xr-robust) through fraction allelic expression analysis (Fig. 5C). Allelic X:A ratio analysis showed not much change in active-X:A ratio in sgXIST, inhibitor-treated and sgXIST+inhibitor cells compared to the WT cells for the reactivated genes cohort (Fig. 5D). Allelic expression of X-linked genes compared to the autosomal genes also showed that there was still higher active-X expression compared to the autosomal allelic expression in sgXIST, inhibitor-treated and sgXIST+inhibitor cells (Fig. 5E). Only, inhibitor treated cells showed a decrease in allelic active-X:A ratio indicating slight erasure of X-upregulation. Finally, gene-wise allelic expression analysis revealed little change in their expression from the active-X chromosome allele in sgXIST and sgXIST+inhibitor cells (Fig. 5F). However, few genes in inhibitor-treated cells showed a slight reduction from active X expression compared to the control B-cells. Overall, it appeared that in B-cells, partial reactivation of inactive-X genes does not trigger the erasure of X-upregulation globally nor from their counterparts on the active X chromosome.

**Figure 5:**
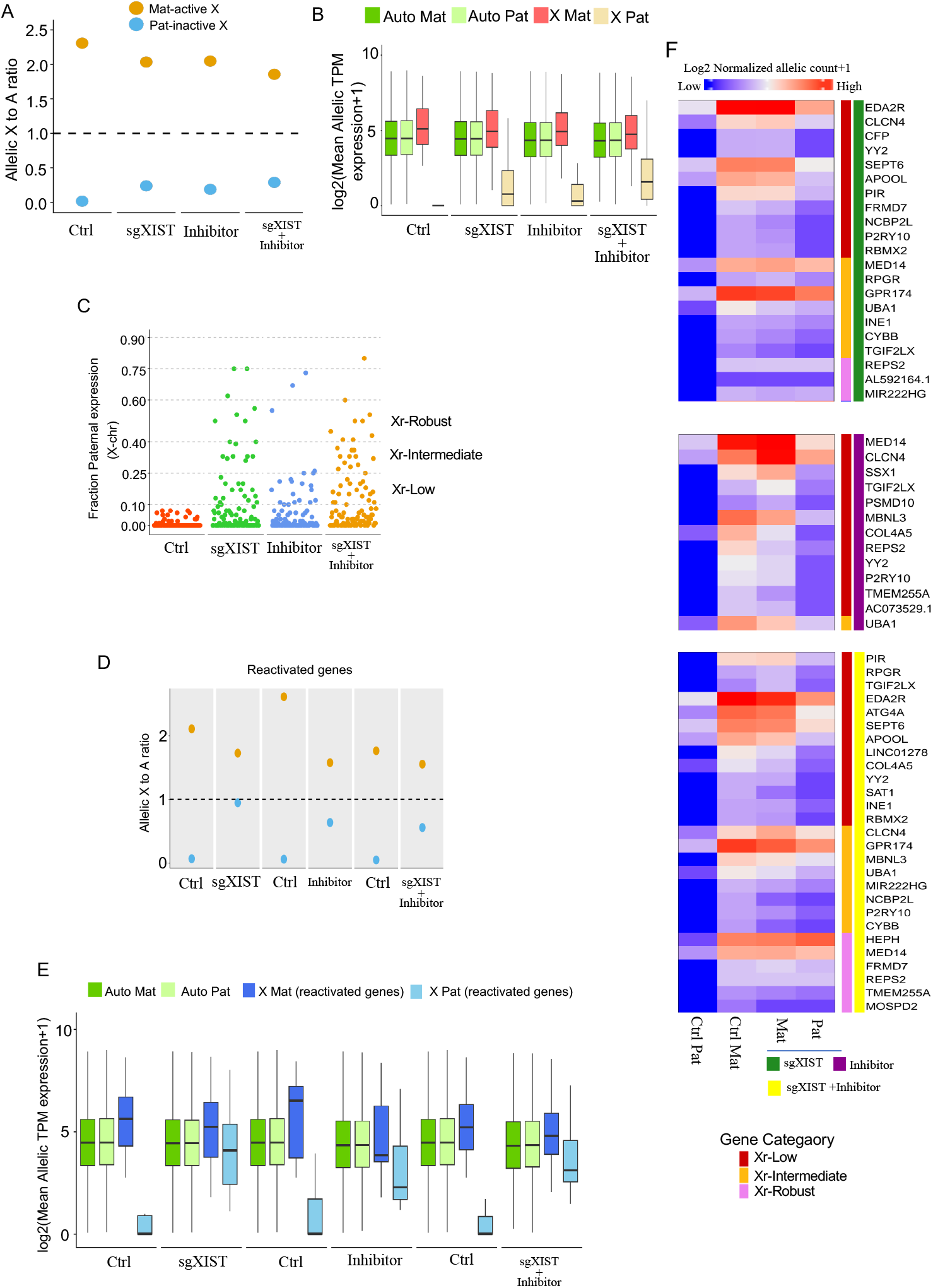
Partial reactivation of inactive X in B cells does not lead to the erasure of X-upregulation. Plots representing (A) allelic X:A ratio (B) autosomal and X-chromosome allelic expression (log mean allelic TPM+1) in Ctrl and sgXIST, inhibitor and sgXIST + inhibitor treated B-cells. (C) Fraction paternal X-chromosome expression in Ctrl and sgXIST, inhibitor and sgXIST + inhibitor treated B-cells. Plots for (D) allelic X:A ratio for reactivated X-linked genes and (E) allelic expression (log mean allelic TPM+1) plot for autosomal and reactivated X-linked genes. (F) Heatmap representing allelic expression (Log2 normalized allelic reads) of X-linked genes from Xi-Pat and Xa-Mat allele in Ctrl and sgXIST, inhibitor and sgXIST + inhibitor treated B-cells.

### Putative regulators of erasure of active X upregulation

Factors and pathways involved in the erasure of X-upregulation are not known yet. We thought that we could use the variability of the transcriptome of cells undergoing erasure of X-upregulation vs. not erasing X-upregulation to identify putative regulators of erasure of X-upregulation. Therefore, we performed a comparative analysis between cells having erasure of X-upregulation (XaXa-E5.5 EPI/iPSC) and cells without erasure of upregulation (XuXa-germ cells) upon X-reactivation (Fig. 6A). First, we profiled differentially expressed genes between XaXa-E5.5 EPI/iPSC and XuXa-germ cells (Fig. 6A and 6B). Next, we focused on differentially expressed transcription factors (TFs) having direct target sites to X-chromosome, which might be the potential candidates for the erasure of X-upregulation. From the differentially expressed genes, we filtered out 25 TFs having > 50 X-linked loci binding sites as estimated from the https://tflink.net/ database, which has well-annotated information regarding TF and its target (Fig. 6C and 6D). Among these 25 TFs, 3 TFs (Sumo2, Smarca5 and Stra8) showed higher expression in XuXa-germ cells than in the XaXa-E5.5 EPI/iPSC (Fig. 6D). We propose that Sumo2, Smarca5 and Stra8 may play an inhibitory role towards the erasure of X-upregulation in XuXa-germ cells. On the other hand, the other 22 TFs showed higher expression in XaXa-E5.5 EPI/iPSC cells compared to the XuXa-germ cells (Fig. 6D). Further, we explored the expression dynamics of these 22 TFs during iPSC reprogramming and if their expression can be correlated with the of erasure of X-upregulation. To find this, we projected the expression pattern of these 22 TFs at the single-cell level to the XuXi cells and XaXa cells throughout the reprogramming (Fig.6E, Fig.S3). Similarly, we profiled the expression of these TFs in XuXi and XaXa embryonic epiblast cells (Fig.6E, Fig. S3). We found that 5 TFs (UTF1, MYCN, DNMT3A, CBX7 and USF1) had very high expression in X-upregulation erased XaXa cells compared to the XuXi cells in both iPSC and embryonic epiblast cells (Fig.6E, Fig. S3). Taken together, we propose that these five TFs are the potential driver of erasure of X-upregulation in XaXa-E5.5 EPI/iPSC cells. As XuXa-germ cells lack these TFs, they are unable to erase X-upregulation. In the future, further experiments to be conducted to elucidate the role of these factors in the erasure of X-upregulation.

**Figure 6:**
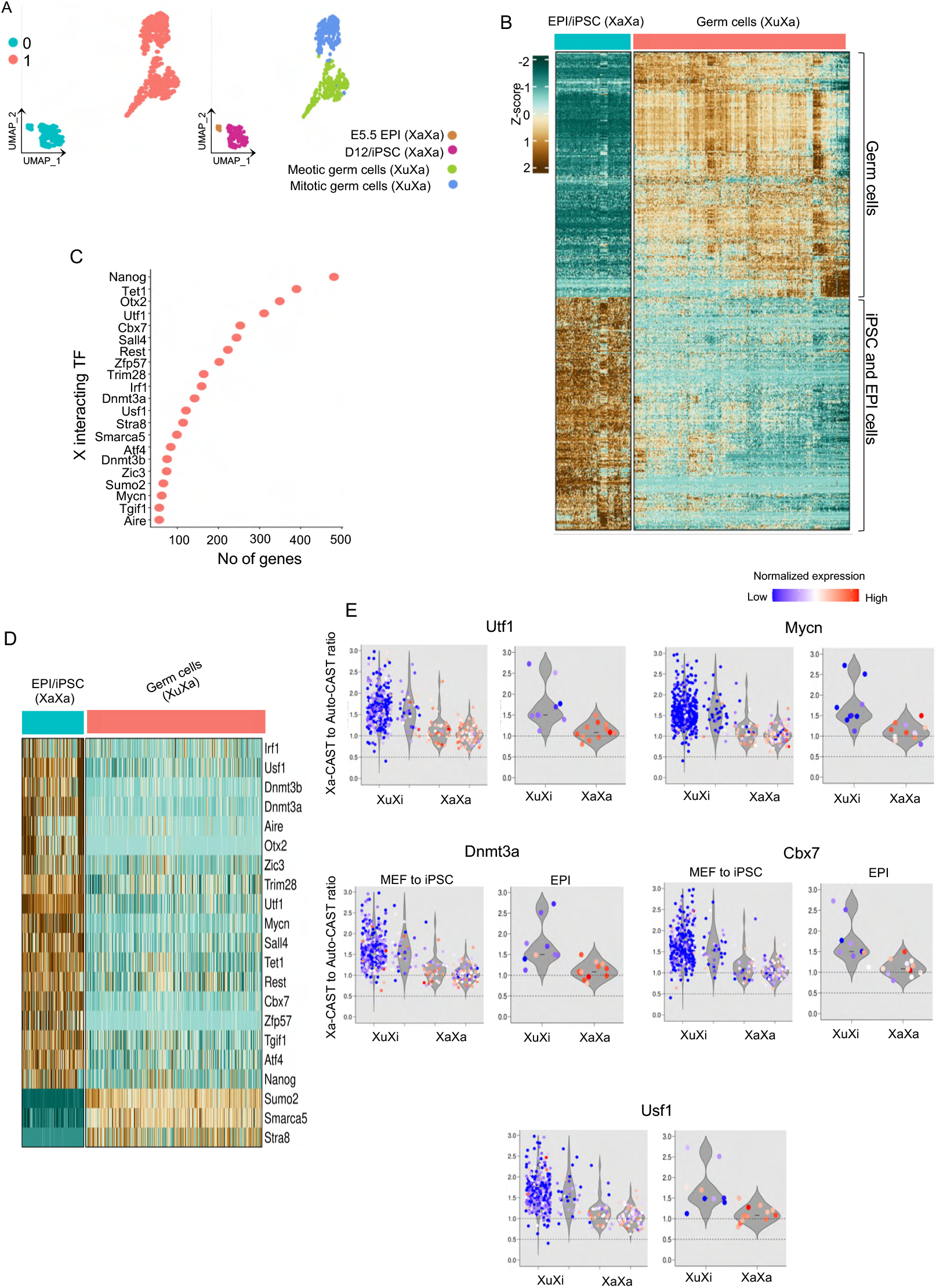
Identification of putative regulators of erasure of X-upregulation. (A) UMAP clustering of XaXa-E5.5 EPI/iPSC and XuXa-germ cells. (B) Heatmap representing differentially expressed genes between XaXa-E5.5 EPI/iPSC and XuXa - germ cells. (C) Plot representing TFs having > 50 X-linked loci binding sites. (D) Heatmap showing the differential expression of TFs having > 50 X-linked loci binding sites between XaXa-E5.5 EPI/iPSC and XuXa - germ cells. (E) Violin plots projecting the expression pattern of TFs (UTF1, MYCN, DNMT3A, CBX7, and USF1) at the single-cell level to the XuXi and XaXa cells throughout the reprogramming of female MEF to iPSC and embryonic epiblast cells.

### Self-inhibition with cross-inhibitory regulation explains X-chromosome dynamics during X-reactivation

Next, we attempted to establish a phenomenological mathematical model to understand the nature of interactions between two X-chromosomes during X-chromosome reactivation. Our experimental data have shown dynamic erasure of X-upregulation upon reactivation of the inactive-X during iPSC reprogramming. We simulated different alternative models and asked what nature of interactions between the two X-chromosomes best explain the observed X-chromosome dynamics during iPSC reprogramming based on mean allelic X:A ratio at different stages of reprogramming at the population level (Fig. 3). We also tested different models in the context of partial reactivation as observed in sgXist XEN cells (Figure 4). In this case, due to the lack of temporal data, we assumed a hypothetical case of iPSC reactivation stalling at day 12 and the value at day 12 was extrapolated up to day 15. These values match qualitatively with the partial reactivation state of sgXist XEN cells. In our modeling framework, each X chromosome was considered a single entity, and we used the X:A ratio as the activity level for that particular X-chromosome.

First, we verified the fits related to X:A ratio dynamics of the active and inactive-X chromosomes during iPSC reprogramming using antagonistic/inhibitory cross-regulation between the two chromosomes. We obtained poor fits, as represented in Fig. 7A, suggestive of the fact that the model must be modified. Next, we added activatory self-regulations for each chromosome. The fits obtained by the addition of self-activations were much better than those without self-activations (Fig. 7B). Similarly, the fits obtained by the addition of self-inhibition were also satisfactory (Fig. 7C). Taken together, this analysis indicated that some form of self-regulation is necessary to explain the observed X-chromosome dynamics. Next, we considered all combinations of cross-regulatory links (interactions between the chromosomes) and self-regulatory links (interactions within a chromosome) (Fig. 7D). We first tested the self-regulatory connections while the cross-regulatory connections were kept fixed as inhibitory. For the full reactivation case, we observed that the connection to Xa being inhibitory gives a good fit regardless of the connection to Xi (Fig. 7E). The case where both were self-activatory also performed well. However, for the case of partial reactivation (Fig. 7F and 7G), most of these fail, and only the case where both are self-inhibitory performs the best. Next, we tested the cross-regulatory connections while fixing the self-regulatory connection as inhibitory. Here, we observed that the case that best fits the full reactivation case (Fig. 7H) is when both the cross-connections are inhibitory. Whereas, for the partial reactivation case (Fig. 7I), the incoming connection to Xa being inhibitory gives a better fit, while the connection to Xi being activatory performs slightly better. Similarly, we also tested the cross-regulatory connections while fixing the self-regulatory connection to be activatory. We found that the incoming connection to Xi does not matter as much for the full reactivation case (Fig. 7J). However, the incoming connection to Xa being inhibitory gives a better fit (Fig. 7J). Similarly, the incoming connection to Xa being inhibitory for the partial reactivation case (Fig. 7K) provides a better fit, whereas the incoming connection to Xi does not matter as much. Altogether, our simulation results indicate that the self-inhibition with cross-inhibitory regulation better explains the X-chromosome dynamics during reactivation of the inactive-X consistently for both partial and full reactivation.

**Figure 7:**
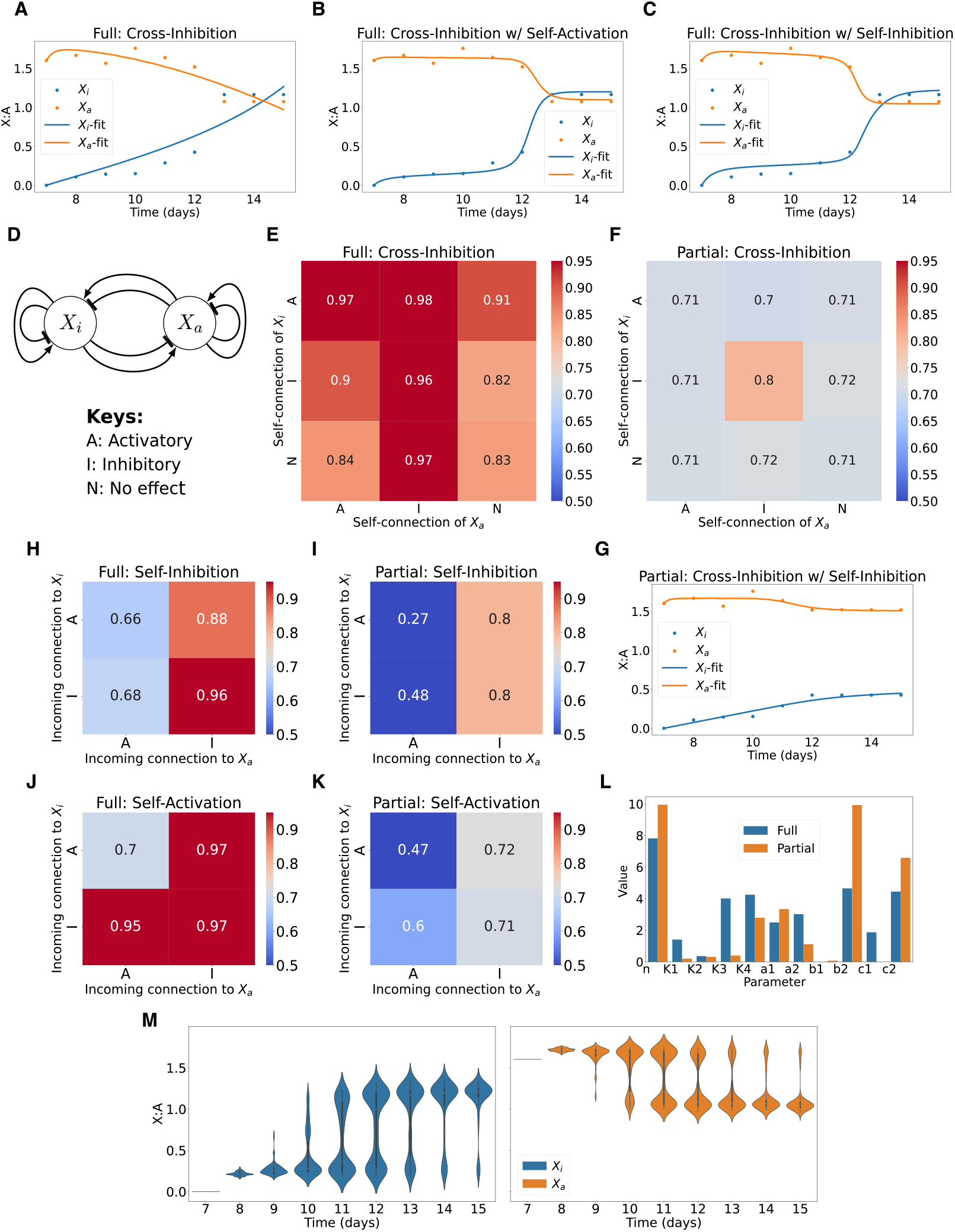
Phenomenological model to explain partial and full reactivation dynamics. Plots representing fits obtained from simulations related to X:A ratio of two X chromosomes during iPSC reprogramming: (A) with only cross-inhibition, (B) with cross-inhibition and self-activation, and (C) with cross-inhibition and self-inhibition. (D) Schematic representation of all combinations of possible cross-regulatory links (interactions between the chromosomes) and self-regulatory links (interactions within a chromosome). (E) and (F) Heatmap representing *R* for different fits for testing self-regulatory connections with fixed cross-inhibition on full and partial reactivation data, respectively. (G) Plot representing fits on allelic X:A ratio during partial reactivation with crossinhibition and self-inhibition. (H) and (I) Heatmaps of *R*^2^ for fits for testing cross-regulatory connections with fixed self-inhibition on full and partial reactivation data, respectively. (J) and (K) Heatmaps of *R*^2^ for fits testing cross-regulatory connections with fixed self-activation on full and partial reactivation data, respectively. (L) Comparison of different fit parameters with cross-inhibition and self-inhibition between full and partial reactivation. (M) Time-course distribution of X level on addition of noise to the model.

Finally, to identify the difference between the two cases (full vs. partial reactivation), we quantified the coefficients of each term, which can act as a proxy for the strength of connections (Fig. 7L). The cross-inhibition on Xi given by a1 was higher in the case of partial reactivation, whereas the cross-inhibition on Xa given by a2 was lower in the case of partial reactivation. The self-inhibition on Xa given by b2 was higher in the case of partial reactivation, whereas the self-inhibition on Xi is negligible in both cases (Fig. 7L).

Our model so far has been fit to the mean X:A values at the population level. However, we observe heterogeneity in the population in the experimental data, i.e., at any time point, there is a fraction of the population having inactive-X (XaXi) and the others which initiated reactivation of the inactive-X (Xa Xa). Furthermore, this fraction keeps changing over time. We hypothesized that the heterogeneity can be explained by the presence of noise in the system. To test this hypothesis, we added a noise term to our equations and solved them with the parameter set we obtained from our fits. We see here that the addition of noise leads to a fraction of cells reactivating faster than the others in the system, evident by the bimodality (Figure 7M). Hence, our model can also explain the heterogeneity in reactivation in addition to the mean behavior of the population.

## Discussion

X-linked genes play a vital role in different biological pathways, including early development, cell fate specification and pluripotency exit (Cloutier et al., 2022; Marahrens et al., 1997; Schulz et al., 2014; Zhang et al., 2016). However, X-linked genes in male and female cells are expressed monoallelically. Therefore, maintenance of appropriate dosage of X-linked gene expression relative to autosomal genes in different developmental contexts is crucial. Recently, studies by us and others have shown that the upregulation of active-X chromosome both in male and female cells during early development fine-tunes the X-linked gene expression as proposed by Ohno (Lentini et al., 2022; Mandal et al., 2020; Naik et al., 2022). Importantly, we demonstrated that during the initiation of random X-inactivation in female pre-gastrulation embryos, X-upregulation happens in a highly coordinated manner along with the X-inactivation (Naik et al., 2022). Here, we have profiled coordination between two X-chromosomes in female cells in different developmental contexts. Like the random X-inactivation, we found that initiation of imprinted X-inactivation in mouse pre-implantation embryos is linked with concomitant upregulation from the active X chromosome (Fig. 1). Our result is consistent with a recent study by Lentini et al. (Lentini et al., 2022). Altogether, it appears that two X-chromosomes in female cells coordinate with each other to fine-tune the X-linked gene expression during early development. Similarly, we show that active X-chromosome in male cells also undergo upregulation during early embryogenesis (Fig. 1). Taken together, our study infers that X-upregulation is integral to the X-linked gene expression homeostasis.

Next, we investigated if there is such coordination between the two X-chromosomes during inactive-X reactivation in different developmental contexts. Interestingly, we found that the reactivation of inactive-X in the epiblast cells of the mouse embryo is associated with the erasure of X-upregulation (Fig. 1I, K). On the other hand, there was slight erasure of X-upregulation in male cells at the similar time point of embryonic development (Fig. 1J). Similarly, we found dynamic erasure of active-X upregulation upon reactivation of inactive X during iPSC reprogramming (Fig. 2). Surprisingly, we do not see such erasure of X-upregulation upon reactivation of the inactive-X during differentiation of PGCLCs toward meiotic germ cells, suggesting that reactivation of the inactive X is not always linked to the erasure of X-upregulation instead it may occur in a lineage-specific manner (Fig. 3). We acknowledge that our observation is based on an *in vitro* system that mimics the meiotic entry of PGCs, therefore in the future studying X-chromosome dynamics *in vivo* would validate further our observation. Indeed, previous studies demonstrated that the erasure of X-upregulation in germ cells is not linked with the X-reactivation and might occur in later stages (Sangrithi et al., 2017). Moreover, the appropriate X-chromosome dosage is believed to be necessary for PGC maturation. It was shown that XO or XY germ cells are not potent like XX germ cells for meiotic initiation (Hamada et al., 2020). On the other hand, Severino et al. have shown how the state of X-inactivation and reactivation during PGC maturation is crucial for efficient meiotic entry (Severino et al., 2022). They found that PGCLCs which underwent robust reactivation of the inactive-X have less potential for meiotic differentiation. Our analysis revealed that cells still in mitotic state have hyperactivation of the inactive-X and active-X upregulation is still maintained. Therefore, it may be possible that hyperactivation of inactive-X coupled with no erasure of X-upregulation in these mitotic cells creates an extra X-chromosome dosage, thereby limiting their potential towards meiotic entry (Fig. 3H). On the other hand, although meiotic cells have still upregulated X, they do not have hyperactivation of the inactive X, which may have favored their meiotic entry (Fig. 3H). Additionally, it may be possible that maintaining active-X upregulation is a prerequisite for meiotic entry. Indeed, in an *in vivo* study, it has been reported that E12.5 to E14.5 germ cells have an excess dosage of X-linked gene expression just before meiotic entry (Sangrithi et al., 2017). In the future, more extensive studies are necessary to understand the relevance of X-chromosome dynamics to PGC maturation. Nevertheless, our study using *in vitro* germ cells establishes the fact that reactivation of the inactive-X always does not trigger the erasure of X-upregulation.

On the other hand, we show that in sgXist XEN cells, partial reactivation of X-linked genes does not trigger the erasure of X-upregulation, indicating that extensive chromosome-wide reactivation is might necessary to trigger the erasure of X-upregulation (Fig. 4). Moreover, we show that even the counterparts of reactivated X-linked genes on the active-X do not undergo erasure indicating that erasure upon reactivation may not occur in a gene-wise manner (Fig. 4). We had an almost similar observation in human B cells where there was partial reactivation of inactive X, but not much erasure of X-upregulation (Fig. 5).

Next, we have identified five TFs: UTF1, MYCN, DNMT3A, CBX7 and USF1 as putative regulators for the erasure of X-upregulation (Fig. 6). We found that the expression of all these TFs increases dramatically in cells with X-upregulation erasure (XaXa) compared to the XuXi cells during iPSC reprogramming (Fig. 6E). Similarly, in the embryonic epiblast, XaXa cells that underwent erasure of X-upregulation have very high expression of these TFs compare to the cells harboring upregulated X (XuXi). Moreover, all these factors are lowly expressed in late germ cells. Taken together, we propose that UTF1, MYCN, DNMT3A, CBX7 and USF1 mediated erasure of X-upregulation during iPSC reprogramming and in the embryonic epiblast. Since germ cells lack these factors, they are unable to erase X-upregulation. Among these factors, we believe that UTF1 is one of the most promising candidates for the following reasons. Firstly, UTF1 is a chromatin-associated transcriptional repressor protein with histone-like properties and plays an important role in chromatin organization (Van Den Boom et al., 2007). Secondly, during early embryogenesis, the expression of UTF1 surges specifically in ICMs exactly when erasure of X-upregulation happens i.e., early to late blastocyst stage and its expression go down rapidly at later developmental stages (Galonska et al., 2014; Okuda et al., 1998; Raina et al., 2021). Similarly, UTF1 expression is high in ES cells, which undergo erasure of X-upregulation, and rapidly decrease upon differentiation (Van Den Boom et al., 2007). Thirdly, the expression of UTF1 surges during iPSC reprogramming (Fig. 6 X). Indeed, previous reports suggest that Utf1 is one of the crucial gene required for the maturation of iPSC and Utf1 overexpression dramatically enhances the reprogramming efficiency (Zhao et al., 2008). On the other hand, though Utf1 expresses in migrating PGCs, its expression decreases in late PGCs at the onset or undergoing meiosis (Chuva De Sousa Lopes et al., 2005; Ge et al., 2021). Notably, a previous study demonstrated that UTF1 binds to the X-chromosome in ES cells and depletion of UTF1 in ES cells leads to the upregulation of many X-linked genes suggesting its role in the maintenance of X-linked gene expression (Kooistra et al., 2010). Altogether, UTF1 is a strong candidate to be involved in the erasure of X-upregulation. On the other hand, DNMT3A, CBX7 and USF1 can also play major role in erasing X-upregulation. Specially, polycomb repressive complex 1 (PRC1) protein CBX7 and DNMT3A are shown to form complex and act together in repressing gene expression (Mohammad et al., 2009). Similarly, USF1 has been shown to mediate the recruitment of PRC1 complex (Scelfo et al., 2019). Therefore, it is highly possible that USF1, CBX7 and DNMT3A act together to facilitate the erasure of X-upregulation. In the future, more extensive experimental analysis can validate more about the role of these TFs in X-upregulation pathways.

Separately, through different alternative simulation approaches, we identified that self-inhibition and the cross-inhibition between X-chromosomes better explain the observed dynamics of X-chromosomes during iPSC reprogramming. We believe this model will pave the path for future investigation of different molecular networks involved in such interactions. One possibility is that the cross-inhibition can be mediated through different *trans*-acting factors and/or competition between the two chromosomes for different *trans-acting* activators, as we reported previously during initiation of random X-inactivation (Naik et al., 2022). On the other hand, self-inhibition can be mediated through cis-acting repressors (Mutzel et al., 2019). Finally, we think the mechanisms mediating such interactions among the X-chromosomes would be pretty complex.

## Materials and Method

### Data availability

Raw data (RNA-Sequencing) generated in this study will be deposited to GEO and made available to the public after publication. Data can be available by the corresponding author upon request. The previously published dataset used for this study is available at Gene Expression Omnibus under the following accessions: Pre-implantation embryos-GSE45719 (Deng et al., 2014), GSE80810 (Borensztein et al., 2017a), GSE89900 (Borensztein et al., 2017b), GSE74155(Chen et al., 2016), Post-implantation: GSE109071(Cheng et al., 2019), iPSC-GSE153846 (Talon et al., 2021), GSE126229 (Janiszewski et al., 2019), B-cells-GSE164596 (Yu et al., 2021), Germ cells: GSE169201 (Severino et al., 2022).

### Cell culture

XEN cells were cultured using media Dulbecco’s modified eagle medium (DMEM) (Hi-media, #AL007A) supplemented with fetal bovine serum (FBS, Gibco #10270-106) L-glutamine (Gibco #25030081) Non-essential amino acids (NEAA, Gibco #11140050), penicillin-streptomycin (Gibco, # 15140122)1mM of 2-Mercaptanol (Sigma #M6250).

### Ablation of Xist in XEN cells using CRISPR-Cas9

Ablation of Xist in XEN cell was achieved through the conventional CRISPR-Cas9 tool. In brief, we used two small guide RNAs (sgRNAs): sgRNA1:AATCCGGAGTATGAGCTGAG sgRNA2: TGCTGCGAAGGAATTTACAG targetting upstream of Xist and intron 1. sgRNAs were designed using CHOPCHOP (https://chopchop.cbu.uib.no). The sgRNAs were cloned into the pSpCas9(BB)-2A-Puro plasmid (PX459) and transfected to XEN cells using lipofectamine 2000 (Invitrogen, # 11668030). After selection through puromycin (3μg/ml) (Takara #631305), colonies were picked up and expanded for screening through Xist RNA-FISH.

### RNA fluorescence in situ hybridization (RNA-FISH)

We generated double-stranded RNA-FISH probes as described previously (Gayen et al., 2015). In brief, probes were generated through random priming of BAC DNA using the Bioprime labeling kit (Invitrogen, #18094-011). Probes were labeled with Cy3-dUTP or Cy5-dUTP (Enzo Life Sciences) and purified through ProbeQuant G-50 Micro columns (Cytiva, #28903408). Probes were precipitated using 0.3M sodium acetate (Sigma, #71196), 300 μg of Yeast tRNA (Invitrogen, #15401011), 150 μg of sheared Salmon sperm DNA (Invitrogen, #15632-011) and absolute ethanol (Hayman, #F205220) at 13,000 rpm for 20 mins at 4°C. The pellet was washed with 70% and followed by 100% ethanol. After washing, probes were dried and resuspended in deionized formamide (VWR Life Sciences, #0606), followed by denatured at 95°C. Finally, probes were preserved at −20°C in a hybridization solution containing 20% Dextran sulfate (SRL, #76203), 2X SSC (SRL, #12590).

For RNA-FISH, XEN cells were seeded on the coverslip and grown to ~60-70% confluency and permeabilized with ice-cold cytoskeleton buffer (CSK, 100 mM NaCl, 300 mM sucrose, 3 mM MgCl2, and 10 mM PIPES buffer [pH 6. 8]) with 0.4% triton-X (SRL #30190). Next, cells were fixed through 3% paraformaldehyde solution (PFA Electron Microscopy Sciences #15710) for 10 min and followed by washed three times with ice-cold 70% ethanol. Next, cells were dehydrated through an ethanol series of 70%, 85%, 95% and 100% and subsequently air-dried. Cells were then hybridized with double-stranded probes for overnight at 37°C in a humid chamber. The cell samples were then washed 3× with pre-warmed 2X SSC/50% Formamide, 2X SSC, and 2 times with 1X SSC for 7 mins each at 37°C. DAPI (Invitrogen, #D1306) was added during the third 2X SSC wash. The coverslips were finally mounted using Vectashield (Vector Labs, #H1000) and visualized under the microscope.

### RNA-sequencing and analysis

Total RNA was isolated using TRIzol (Life technologies #15596-026) according to the manufacturer protocol. Approximately 1ug total RNA with RIN value above 7 was used for library preparation. RNA-Seq library was prepared following the manufacturer’s protocol using the NEBNext library preparation kit for Illumina. Libraries were sequenced on Illumina HiSeq 2500 platform using 2 × 150 bp chemistry. Transcriptomic sequencing reads were mapped against mouse genome GRCm38 (mm10), and human genome GRCh37 (hg19) using STAR (2.7.9a) (Dobin et al., 2013) with default parameter and aligned reads were counted using HTSeq-count (2.0.2) (Anders et al., 2015). The expression level of transcripts was calculated using TPM (Transcripts Per Kilobase Million) counts.

### Allele-specific analysis

We performed allele-specific analysis of RNA-Seq data as described previously (Naik et al., 2021, 2022). In brief, we created in silico reference genome by incorporating strain-specific SNPs in to the mm10 reference genome. Strain-specific SNPs obtained from Mouse Genomes Project (https://www.sanger.ac.uk/science/data/mouse-genomes-project). Reads were mapped separately to parental genomes using STAR. For removing any false positives in allelic count, we only considered those SNPs with minimum read counts of 10 (bulk data) and 3 (single-cell data) per SNP site and used at least 2 informative SNPs per gene. We calculated allelic read counts by taking an average of SNP-wise reads. We normalized allelic read counts across cells using scaling factors obtained from DESeq2 using non-allelic count. The allelic ratio was calculated using formula: (Allele-A or Allele-B/ Allele-A + Allele-B). Allelic TPM fraction (A or B) was calculated using formula: Allelic ratio (A or B) * TPM of the gene. Here, Allele-A and Allele-B are respective strains.

### Single cell clustering

To identify different lineages in pre-implantation embryos, we performed dimension reduction UMAP using seurat package (Butler et al., 2018; Stuart et al., 2019) by RunUMAP function with parameters 1000 highly variable genes and dims 1:30. Parameters used for clustering iPSCs:HVG=2000, dims=1:15 and germ cells: HVG=3000, dims=1:40.

To perform differential gene expression between iPSCs/EPI (XaXa) vs. germ-cells (XuXa) using single cell data, first we clustered cells using seurat (HVG=2000, dim 1:20) and thereafter, cluster specific genes were identified using the function FindAllMarkers (min.pct=0.50, only.pos=TRUE, logfc.threshold=1, test.use=“MAST”).

### Allelic X to A ratio

Considering the huge difference in the number of X-linked and autosomal genes, we calculated allelic X:A ratio using bootstrapping procedures as described previously (Naik et al., 2022; Pacini et al., 2021). In brief, we calculated allelic X:A ratio by dividing the allelic expression of X-linked genes with the allelic expression of the same number of autosomal genes selected randomly for each cell/sample. This was repeated 1000 times and median of 1000 values was considered. To exclude low-expressed genes from our analysis, we used genes having >=5 TPM for bulk RNA-Seq data and >=1 TPM for scRNA-Seq data. Escapees and pseudo autosomal genes were not considered for this analysis.

### Sexing of the embryo

Sexing of the available single-cell dataset used for this study was performed if the sex was not mentioned previously. Cells were assigned as male based on Y chromosomal genes expression (Zfy2, Zfy1, Kdm5d, Uty, Usp9y, Ddx3y, Eif2s3y, Ube1y1).

### Simulation

We have considered the two X-chromosomes as interacting entities, and they are modeled as differential equations given by:

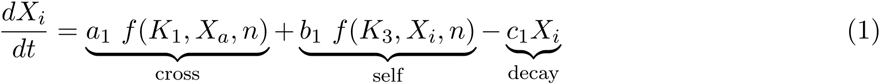

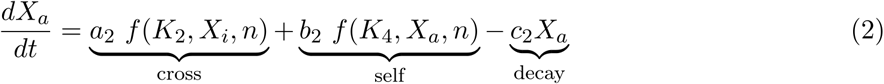

Where,

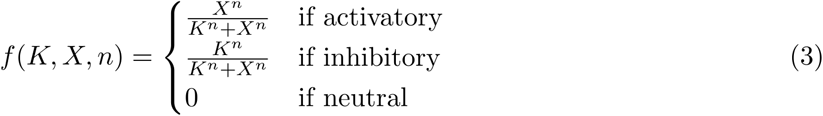

*X_i_* is the expression level of the inactive X given as X:A ratio, *X_a_* is the expression level of the active X given as X:A ratio, *a*_1_,*a*_2_,*b*_1_,*b*_2_,*c*_1_ and *c*_2_ are the coefficients for cross-regulatory, self-regulatory and decay terms, respectively, *n* is the hill coefficient, *K*_1_,*K*_2_,*K*_3_ and *K*_4_ are the half-saturation constants.

For the data for full reactivation, we took the data obtained from X-reactivation in iPSC given in Fig. 2. Due to lack of data between day 0 to day 8, the starting point for X-reactivation was considered to be day 7 and with the same level corresponding to day 0 in the iPSC data. The mean value for day wise levels of *X_i_* and *X_a_* were considered. The levels for iPSC cells were considered as the levels from day 13 to day 15. For the data for partial reactivation, we assumed a hypothetical case of iPSC reactivation stalling at day 12. The value at day 12 was extrapolated up to day 15. These values match qualitatively with the partial reactivation state in Fig. 4 and have been done due to the lack of temporal data.

These equations are fit to the time course data for full and partial reactivation. This was done by minimizing the sum of square error using the differential evolution algorithm of scipy. The initial population of parameters is sampled using Sobol sampling. The differential equations are solved using the explicit Runge-Kutta method of order 5(4) with these parameters. Then, the sum of square errors between the solutions evaluated at the given time points and the actual data is calculated. A new parameter set is generated by adding a weighted difference between two randomly chosen parameter sets to a third parameter set, similar to a mutation. Then, it randomly combines parameters from the old set with this new set, similar to crossover. The sum of square errors with this new set of parameters is also evaluated and compared with those of the old parameters. If the values are lower with the new set, they replace the old set in the next generation of the population. This is repeated multiple times until an optimal solution is found (Storn and Price, 1997).

For the model with noise, we added a noise term *η*(*t*) to Equation 1 and Equation 2 where the values are sampled from a normal distribution with mean 0 and standard deviation 1. These were then solved with the fit parameters using explicit Runge-Kutta method of order 5(4).

## Code availability

The code implementing this is available at https://github.com/Harshavardhan-BV/rev-XCI.

## Visualization and Plots

All plots were generated using R version 4.2.1 using ggplot2 library and Integrative Genomics Viewer (IGV) for genome visualization.

## Author’s Contribution

SG conceptualized, supervised and acquired the funding for the study. HCN, DC, SM, MA, RB, Parichitran and Avinchal performed experiments. MKJ, HBV and KH performed simulation. SG, HCN, SM, MA, MKJ, HBV and KH wrote, edited, and proofread the manuscript. The final manuscript was edited and approved by all the authors.

## Acknowledgments

We thank Prof. Sundeep Kalantry, University of Michigan for providing BACs and cell line. This study is supported by DBT grant (BT/PR30399/BRB/10/1746/2018), DST-SERB (CRG/2019/003067), DBT-Ramalingaswamy fellowship (BT/RLF/Re-entry/05/2016) and Infosys Young Investigator grant award to SG. MA and DC acknowledge Indian Institute of Science (IISc), Bangalore for the fellowship. SM and Avinchal acknowledge University of Grant Commision (UGC) for the fellowship. RB would like to acknowledge council of scientific and industrial research (CSIR), India for the fellowship. H. BV would like to acknowledge Prime minister research fellowship (PMRF), India.

## Conflict of interest

The authors declare that they have no conflict of interest.

## Notes

### Competing Interest Statement

The authors have declared no competing interest.

### Summary of Updates

The manuscript has been extensively revised with new experimental data.

