## Supplementary figures for "Lineage-specific dynamics of erasure of X-upregulation during inactive-X reactivation"

Fig.S1

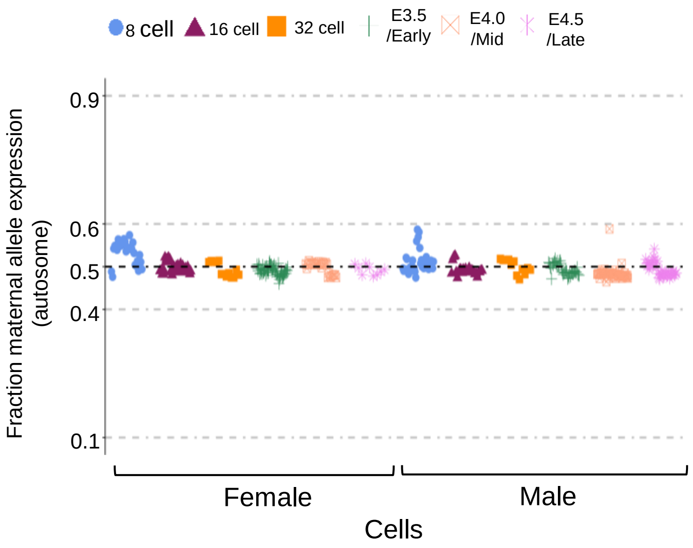
8 cell
16 cell
32 cell
E3.5 /Early
E4.0 /Mid
E4.5 /Late
Fraction maternal allele expression (autosome)
0.9
0.6
0.5
0.4
0.1
Female
Male
Cells

**Figure S1: Allelic expression of autosomal genes in pre-implantation embryos (related to Fig. 1).** Fraction maternal expression of autosomal genes in different stages of mouse pre-implantation hybrid embryos (8-cell, 16 cell, 32-cell, E3.5 early, E4.0 mid and E4.5 late blastocyst).

Fig.S2

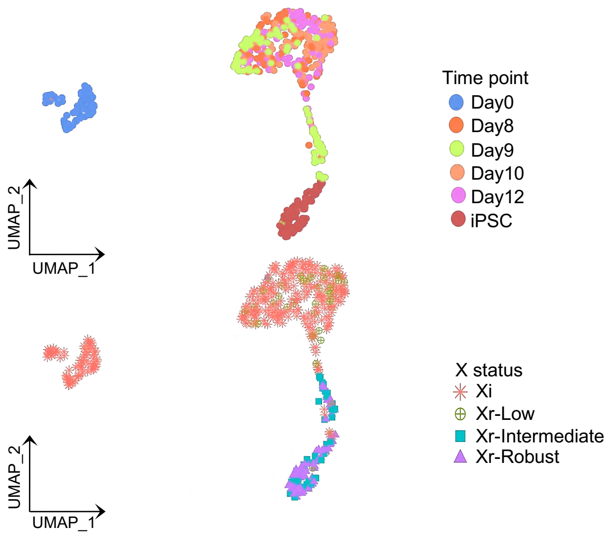
Time point
Day0
Day8
Day9
Day10
Day12
iPSC
UMAP_2
UMAP_1
X status
Xi
Xr-Low
Xr-Intermediate
Xr-Robust
UMAP_2
UMAP_1

**Figure S2: (related to Fig. 2).** Projection of different categories of cells: Xi, Xr-Low, Xr-intermediate and Xr-robust on UMAP clusters of different stages of reprogramming of female MEF to iPSC.

Fig.S3

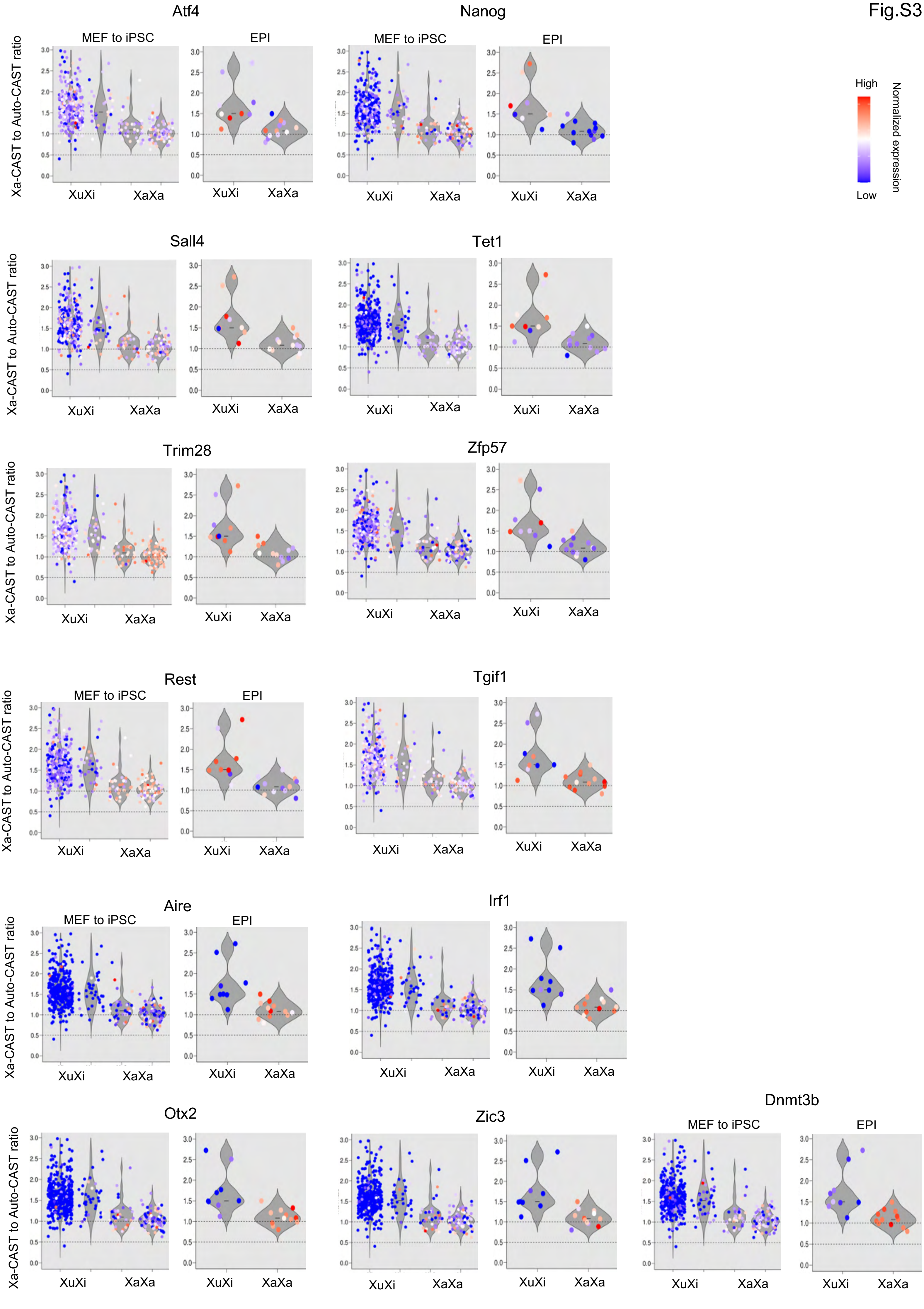

Atf4
Nanog
Sall4
Tet1
Trim28
Zfp57
Rest
Tgif1
Aire
Irf1
Otx2
Zic3
Dnmt3b
MEF to iPSC
EPI
Xa-CAST to Auto-CAST ratio
XuXi
XaXa
3.0
2.5
2.0
1.5
1.0
0.5
0.0
High
Low
Normalized expression

**Figure S3: Expression pattern of TFs during iPSC reprogramming and embryonic epiblast cells (related to Fig. 6).** Violin plots projecting the expression pattern of different TFs at single cell level to the X2aXi and XaXa cells throughout the reprogramming of female MEF to iPSC and embryonic epiblast cells.
